# Electron Videography of a Lipid–Protein Tango

**DOI:** 10.1101/2022.11.13.516315

**Authors:** John W. Smith, Lauren N. Carnevale, Aditi Das, Qian Chen

**Affiliations:** Department of Materials Science and Engineering, University of Illinois at Urbana–Champaign, Urbana, IL 61801, United States; Department of Biochemistry, University of Illinois at Urbana–Champaign, Urbana, IL 61801, United States; Department of Comparative Biosciences, University of Illinois at Urbana–Champaign, Urbana, IL 61801, United States; Beckman Institute for Advanced Science and Technology, University of Illinois at Urbana– Champaign, Urbana, IL 61801, United States; Cancer Center at Illinois, University of Illinois at Urbana–Champaign, Urbana, IL 61801, United States; Department of Chemistry, University of Illinois at Urbana–Champaign, Urbana, IL 61801, United States; Materials Research Laboratory, University of Illinois at Urbana–Champaign, Urbana, IL 61801, United States

## Abstract

Biological phenomena ranging from enzymatic catalysis to synaptic transmission find their origins in the morphological fluctuations of biomolecules and biomolecular assemblies in liquid water. However, directly imaging these nanoscopic dynamics without probes or labels has been a fundamental methodological challenge. Here we developed an approach for ‘molecular videography’—combining liquid-phase transmission electron microscopy with molecular modelling—with which we *filmed* the nanoscale morphological fluctuations of individual, freely suspended, and unlabelled membrane protein nanodiscs in liquid. These unprecedented observations enabled us to quantify fingerprints of nanoscale lipid–protein interactions directly, revealing that nanodisc shape undulations are defined by the molecular mechanics of encircling scaffold proteins, whereas lipid interactions with a membrane protein reach across surprisingly long ranges to delineate dynamically modified membrane domains and increase nanodisc stability. Overall, this work illustrates an experimental approach to image, quantify, and understand biomolecular dynamics at the nanometre scale.

## Main Text

Imagine a *Noh* drama, a flamenco dance, or one of the great silent films. In these art forms without words, form, gestures, and motion tell the story, revealing the traits and relationships of the characters. Though occurring on a smaller stage, the motions of biomolecules do the same. As they undergo structural transformations in native or perturbed states, proteins, lipids, and nucleic acids recount the tales of enzymatic catalysis, synaptic transmission, DNA replication, *etc*. Biological processes are built upon these “dances”^1^, with structural and dynamical features defined by liquid water^2^. Water defines the hydrophobic collapse in protein folding and smooths energy barriers on the route to functional structures, but it also facilitates long-range communication between molecules or subunits, and may enhance or suppress conformational flexibility, in which cases its own structural and dynamical properties enter into biomolecular function^2,3^. As such, fully understanding biomolecular interactions and functions in context requires an interrogation of both structure and structural dynamics, at the individual level and in liquid, but meeting *all* these criteria is technically challenging. While spectroscopy- or scattering-based approaches can reveal structural and dynamical traits simultaneously and with high spatiotemporal resolution^4–6^, their level of insight is limited to whatever information can survive an ensemble averaging process. On the other hand, decades of effort in microscopy have been devoted to filming *single* biomolecules in action^7–10^, but optical methods^10,11^, including super-resolution imaging techniques using labels, still find it challenging to reveal sub-molecular scale structural information. Approaches based on electron microscopy, meanwhile, have the requisite nanometre (or better) spatial resolution^12,13^, but in electron microscopy the crux of technical challenges has been an incompatibility with liquid water, limiting its scope to the study of fixed or frozen (*i*.*e*., immobilized) specimens. Single-particle reconstruction methods can sometimes infer dynamical information indirectly, but the flexible regions of biomolecules can be lost to averaging. These approaches also demand high molecular uniformity—and therefore laborious purification of just one or a few structural states— which may come at the cost of meaningful molecular individuality.

Here we present the first molecular videography of freely suspended biomolecular assemblies in liquid buffer, with liquid-phase transmission electron microscopy (TEM) (**Fig. 1A–E, Supplementary Fig. 1**). This technique is now growing steadily in the materials science community^14–17^, but efforts to film biomolecules have been confounded by their low stability and visibility. Previous work has achieved snapshots of viruses^18^, structural proteins^19,20^, and cells^21^, and in the process helped develop methods for sample preparation and imaging protocols to mitigate electron beam damage^20,22^, but continuous protein dynamics have not yet been observed. With careful experimental adaptations of liquid encapsulation methods, electron dose rates, and total doses (**Supplementary Notes 1–3**), here we achieve real-time imaging for highly extended periods, with morphology sampling sufficient to pair our real-time observations with molecular dynamics (MD) simulations^23^ and biophysical theory^24^, and capture rare events. Surprisingly, our systematic comparisons with available biochemical and biophysical data described below suggest that the specimen sustains a sufficiently reduced amount of damage from imaging electrons— potentially due to graphene encapsulation (see **Supplementary Note 1** and later discussions)—to yield meaningful insight. Although the spatial resolution is not yet at the level that cryo-EM can achieve by combining thousands of images in single-particle reconstruction and the integrity of all chemical groups may not be completely preserved, the obtained molecular movies reveal the liquid-phase structure of these assemblies individually and in sub-molecular detail without averaging, and make it possible to quantify roles of intermolecular interactions in their dynamics.

**Fig. 1.**
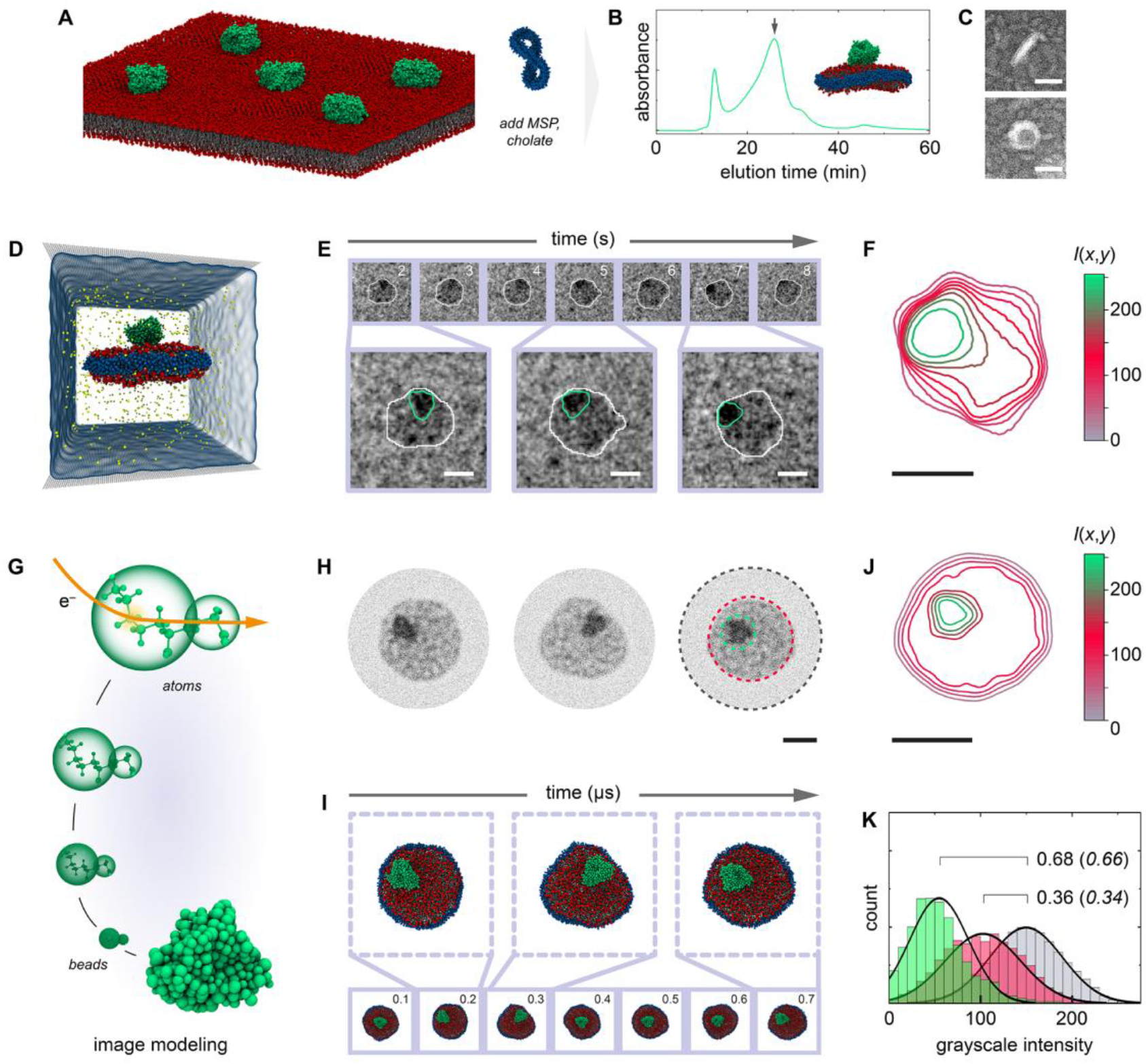
Single-nanodisc imaging by liquid-phase TEM. (**A–B**) Cytochrome P450 2J2 (CYP2J2) was combined with lipids and membrane scaffold proteins (MSPs) to form nanodiscs, with subsequent purification by size exclusion chromatography (fractions at the arrow (**B**) were collected for analysis). In all schemes, CYP2J2 is shown in green, the scaffold proteins in blue, and the lipids shown in red (head groups) and grey (tail groups). (**C**) Negative stain micrographs—where nanodiscs appear bright against the high-atomic number stain background—showing representative elliptical and circular projections. (**D**) Nanodiscs were encapsulated in graphene chambers to image continuous morphology fluctuations using liquid-phase TEM (**E**). (**F**) Contrast variations, highlighted with a representative intensity profile, *I*(*x, y*), reveal the position of CYP2J2 and its orientation in the plane of the membrane (**E**, green lines) within the nanodisc contour (**E**, white lines). (**G–J**) Experimental snapshots were compared with images simulated based on MD-generated coordinates (examples shown in **I**). Contrast variations in simulated images (**J**) matched well with experimental results (**F**). (**K**) The ratio of intensities between the background, lipid-only, and CYP2J2-containing areas (grey, red, and green respectively in **H**, right, and **K**) shows agreement between simulation and experiment (values in regular case are from experimental data, values in italics are from simulation). Scale bars: 20 nm (**C**), 10 nm (**E, F, H, J**).

In this work we focus on how this unique biophysics imaging tool can provide insight into membrane systems, specifically, nanoscale lipid–protein interactions. In particular, we study membrane protein nanodiscs, membrane proteins in a dispersed, “single particle” form now commonly used for investigations of membrane protein structure^25^, but also an ideal model system for studying lipid–protein interactions^26^. Namely, as membrane proteins participate in a wide variety of biological processes—signal transduction, ion transport, membrane metabolism, *etc*.— lipids fine-tune their functionality by dictating the phase behaviour, curvature, and other physicochemical properties of the membrane environment across a range of length and time scales^27,28^. Previous work using nanodiscs using spectroscopy^29,30^ or cryogenic TEM^26^ has shown how membrane proteins are sensitive to local membrane fluidity, and how a few specific, tightly bound lipids can stabilize a protein’s functional state. However, investigating the structure and dynamics of whole nanodiscs—*i*.*e*., including longer length and time scales also relevant to lipid– protein interactions^31^—has remained a challenge. Indeed, nanodiscs also constitute an advantageous test bed for electron videography, as these important thermal fluctuations match the current resolution of the technique. Some understanding has come from MD simulations^26,32^, but these are currently limited to microsecond-scale phenomena. Here we image single-nanodisc structural fluctuations for up to *minute* scales. We achieve a high spatiotemporal resolution (nanometre, sub-second), while preserving complexities that are challenging to model computationally, lending new and quantitative insight into the fundamental molecular motions, mechanics, and lipid–protein interactions that govern nanodisc—and therefore membrane/membrane protein—structural dynamics at the nanometre scale.

## RESULTS

By revealing not only the shape of a nanodisc, but also the extent of its fluctuations, liquid-phase TEM can identify variations in nanodisc size and composition without averaging. For example, we first investigated nanodiscs containing CYP2J2 and stabilized either with a long (MSP 2N2) or short (MSP 1E3D1) scaffold protein (**Fig. 1A–C, Supplementary Figs. 2–3**)^33,34^. CYP2J2 is a cardiac cytochrome P450 involved in arachidonic acid metabolism and a representative peripheral membrane protein whose assembly into nanodiscs is well-established^34^. In graphene-based liquid chambers (**Fig. 1D, Supplementary Figs. 1**,**4–7**), these nanodiscs freely fluctuate and rotationally diffuse (**Supplementary Fig. 8**) for up to a few minutes before succumbing to damage by the electron beam (**Supplementary Fig. 5**), so we monitored their projections over time (**Fig. 1E, Supplementary Video 1**). If projections are fit to an ellipse (**Supplementary Figs. 9–10**), rotation and shape fluctuation manifest in variations of the minor and the major axis, respectively (**Supplementary Note 4**). Nanodiscs with the long MSP display major axes in the range of 18‒22 nm over time, in agreement with the MSP length^35^ and negative stain TEM characterization (**Fig. 1C, Supplementary Fig. 2**). Short MSP nanodiscs are readily distinguishable, with major axes of 13–15 nm (**Supplementary Figs. 2**,**10, Supplementary Table 1**). Meanwhile, empty nanodiscs— prepared with long MSPs but no CYP2J2—had major axes similar to filled ones (∼20 nm) but of a broader statistical distribution (**Supplementary Fig. 10, Supplementary Table 1**). We attribute this broadening to the greater tendency of a nanodisc to fluctuate when hydrophobic or electrostatic interactions between the membrane and a protein are not present. Notably, this also suggests that the influence of a membrane protein on fluctuations reaches the nanodisc boundary—at least several nanometres—which is consistent with MD simulations (**Supplementary Figs. 11–15, Supplementary Tables 1‒2**). Overall, these initial comparative experiments highlight the ability to differentiate nanodiscs structurally, dynamically, and in a robust and quantitative way, with minimized apparent artefacts from the electron beam or substrate (**Supplementary Notes 1–3**). We therefore delved deeper to understand nanodisc properties observed in experiments.

In cases with minor out-of-plane motion (**Supplementary Note 4, Supplementary Figs. 9–10**,**16– 17**), substructural details such as the position and orientation of the membrane protein—as it diffuses and rotates within the nanodisc—can be discerned from intensity variation in nanodisc projections. CYP2J2 is free to explore the interior of the nanodisc and veer “off-center” (**Supplementary Fig. 16**) because the POPC membrane is fluid at room temperature and relatively large compared to the protein itself. Similar behaviour has been proposed based on results from small-angle scattering and cryo-EM^36,37^. Indeed we provide support that roughly triangle-shaped features (**Fig. 1E–F**) have the shape and contrast expected of our membrane protein by comparing experimental TEM images to “synthetic” images simulated based on MD-generated coordinates (**Fig. 1G–I, Supplementary Table 3, Supplementary Video 2**). Simulated images show qualitatively and quantitatively similar contrast variation to experimental video frames, with distinct intensities corresponding to the background, the membrane-only portion of the nanodisc, and the darkest region with the membrane protein (**Fig. 1F,J–K**). This ability to localize and orient proteins in individual lipid–protein assemblies constitutes a major improvement over current experimental tools. Previous imaging work has required nanoparticle labels to monitor membrane protein motion at high resolution^21,38^—an approach which cannot be used to reveal substructural detail and potentially induces abiological fluctuations—or has only revealed topographical information^39^. Our approach may therefore offer opportunities to study positional and orientational dynamics during protein–protein interactions in a membrane environment, such as in docking or oligomer formation.

The evolution of the nanodisc shape over time also offers biophysical insight, about nanodisc interfacial mechanics (**Fig. 2A–B, Supplementary Video 1**). When rotation is negligible, these undulatory motions involve (i) thermal fluctuations of the α-helical MSPs against constraints imposed by hydrogen bonding and (ii) the faster, fluid-like motion of interior lipids (**Fig. 2B–E**). Phenomenologically, undulations decompose into contributions from a range of angular frequencies (**Fig. 2C, Supplementary Fig. 18**). A more physical interpretation of these contributions can be made by analysing the Fourier square amplitude ⟨|*u*(*q*)|^2^⟩ across a range of wavevectors *q* (**Fig. 2F, Supplementary Fig. 19**) ^24^. In doing so, we find that nanodisc contour undulations can be described by two effective mechanical properties: a bending rigidity, *κ*, and a tension, *σ*, with experimental data following 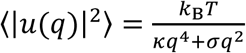 (*k*_*B*_*T* denotes the thermal energy), such that ⟨|*u*(*q*)|^2^⟩ ~ *q*^−2^ in the low *q* regime (with corresponding *σ* = 4.6 × 10^−4^ *k*_*B*_*T* nm^−1^) and ⟨|*u*(*q*)|^2^⟩ ~ *q*^−4^ in the high *q* regime (with corresponding *κ* = 2.1 *k*_*B*_*T* nm). The same regimes appear in equilibrium MD simulations, particularly in the bending-dominated regime (**Fig. 2F, Supplementary Fig. 19, Supplementary Note 3**). We note that these parameters do not seem perturbed by the graphene substrate, with experimental data closely matching “control” simulations where graphene (and beam effects) are not considered (**Supplementary Note 3, Supplementary Fig. 20–21**). Although a comparable bending rigidity has been measured for high density lipoprotein particles—structurally similar assemblies—using nanoindentation^40^, a tension has eluded previous characterization. We anticipate these measures are sensitive to protein content and to the lipids’ phase behaviour, fluidity, or curvature, and offer a metric for future comparative studies. Our simulations show that subtle differences can be detected even for two very closely related peripheral membrane proteins, CYP2J2 and its relative CYP3A4 (**Fig. 2F**). In this system, the interfacial mechanics generally “penalize” deformation, restricting nanodiscs to an overall round shape. Contours are well-described by a single circular domain and local frame-to-frame “flow” (**Fig. 2D–E**) is small and symmetric, with the overall area changing <4%. These mesoscale flows can be directly related to the diffusion of single lipid molecules in MD simulations (**Fig. 2E**).

**Fig. 2.**
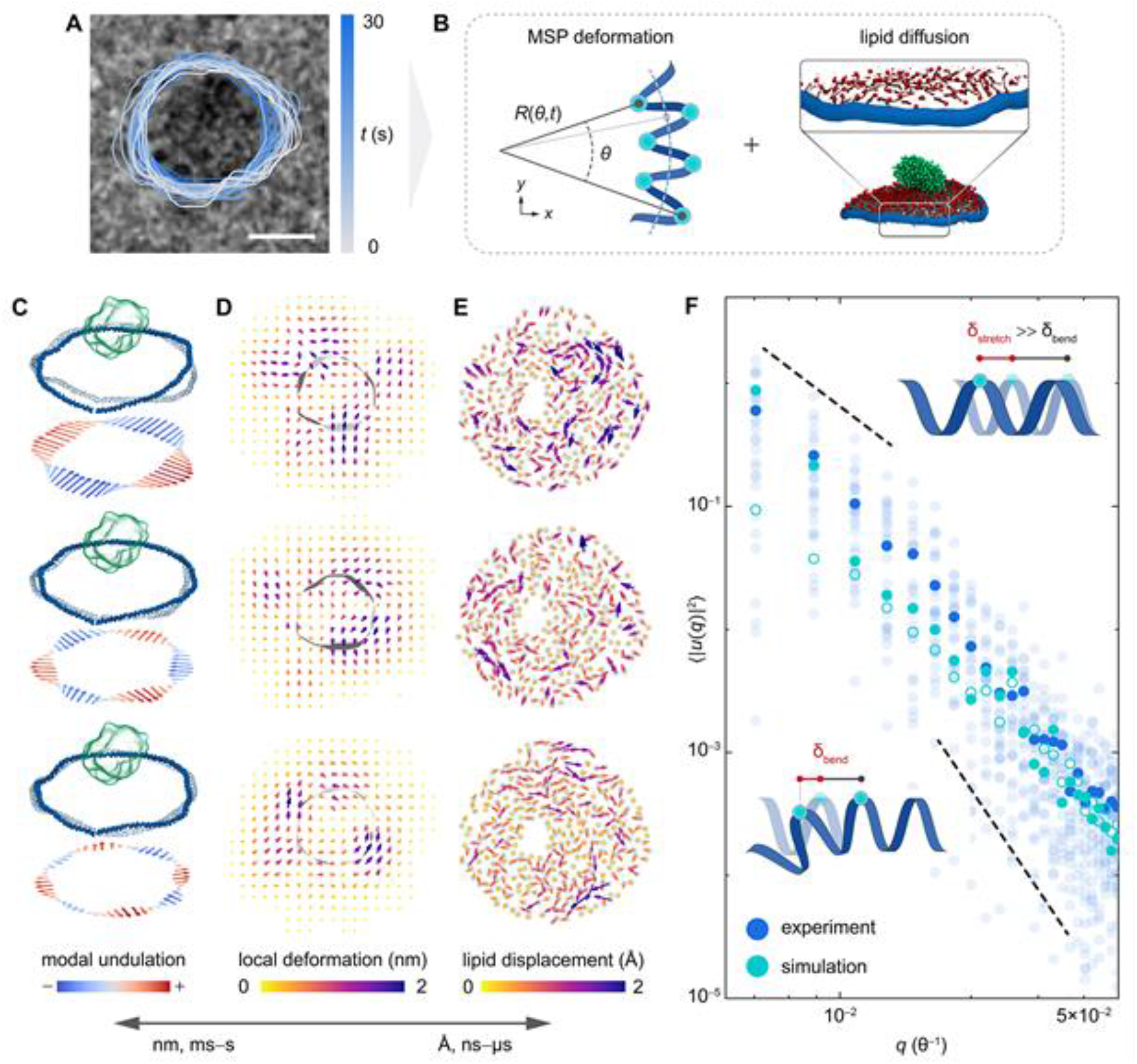
Nanodisc fluctuations as “fingerprints” of molecular phenomena. (**A**) Nanodisc undulations (highlighted with overlaid contours from different time points) involve two components: (**B**) fluctuations of the MSPs, based on a balance between thermal energy and molecular mechanics and fluid-like diffusion of interior lipids. (**C**) Undulations decompose into principally elliptical deformations (56%), with gradually lesser contributions from third-order (22%), fourth-order (6%), and higher-order radial modes (collectively, 16%). These mechanics are coupled to motion of lipids in the nanodisc interior, which can be measured (**D**) on a continuum level, as an overall vector field or (**E**) on the molecular level, following displacements of individual lipid molecules in MD simulations. Individual panels are examples of this “flow” between experimental (**D**) and simulation (**E**) frames. Between experiment and simulation, these measurements span from nanometres and milliseconds to angstroms and nanoseconds. (**F**) Fourier analysis of nanodisc fluctuations suggests that tension dominates at relatively large scales (*δ*_stretch_) whereas bending dominates regimes at smaller scales (*δ*_bend_). The simulation data in solid circles corresponds to results from nanodiscs containing CYP2J2, the data in open circles to those with CYP3A4 (**F**) Scale bars: 10 nm (**A, G**), 20 nm (**F**).

Over the seconds to minute timeframe of our observations, we occasionally captured an unexpected and more extensive form of fluctuation (**Fig. 3A–B, Supplementary Video 3**) in which nanodiscs would “stretch” or “finger” to form distinct domains interdigitated with concave regions (**Fig. 3A–D, Supplementary Figs. 22–23**). These extended deformations were not observed in MD simulations, from the perspective of which they are rare events. We differentiate fingering from the normal undulations based on the contour circularity, a measure of “convexness” (circularities <0.8 were associated with fingering, **Supplementary Fig. 22**). Fingering also converts a nanodisc described by a single, large domain into one comprised of multiple zones approaching 4–5 nm in diameter. Surprisingly, filled nanodiscs that experienced fingering fluctuations eventually regained their overall round shape (**Supplementary Fig. 23, Supplementary Video 3**). When this occurred, the projected area and perimeter of the nanodisc before and after fingering were within ∼8% of each other (**Supplementary Fig. 23a–b**), suggesting minimal damage or mass loss. Fingering nanodiscs also retained a major domain (described by the maximum inscribed circle, with size *d*_*insc*_) of at least ∼12 nm (**Fig. 3C–D**) despite their large overall distortion. In contrast, when empty nanodiscs experienced fingering fluctuations, domains decreased to ∼5 nm before complete disintegration (**Supplementary Fig. 23c, Supplementary Video 3**). Together, these results suggest that (i) filled nanodiscs but not empty ones can recover from fingering fluctuations and (ii) recovery may be facilitated by a central core.

**Fig. 3.**
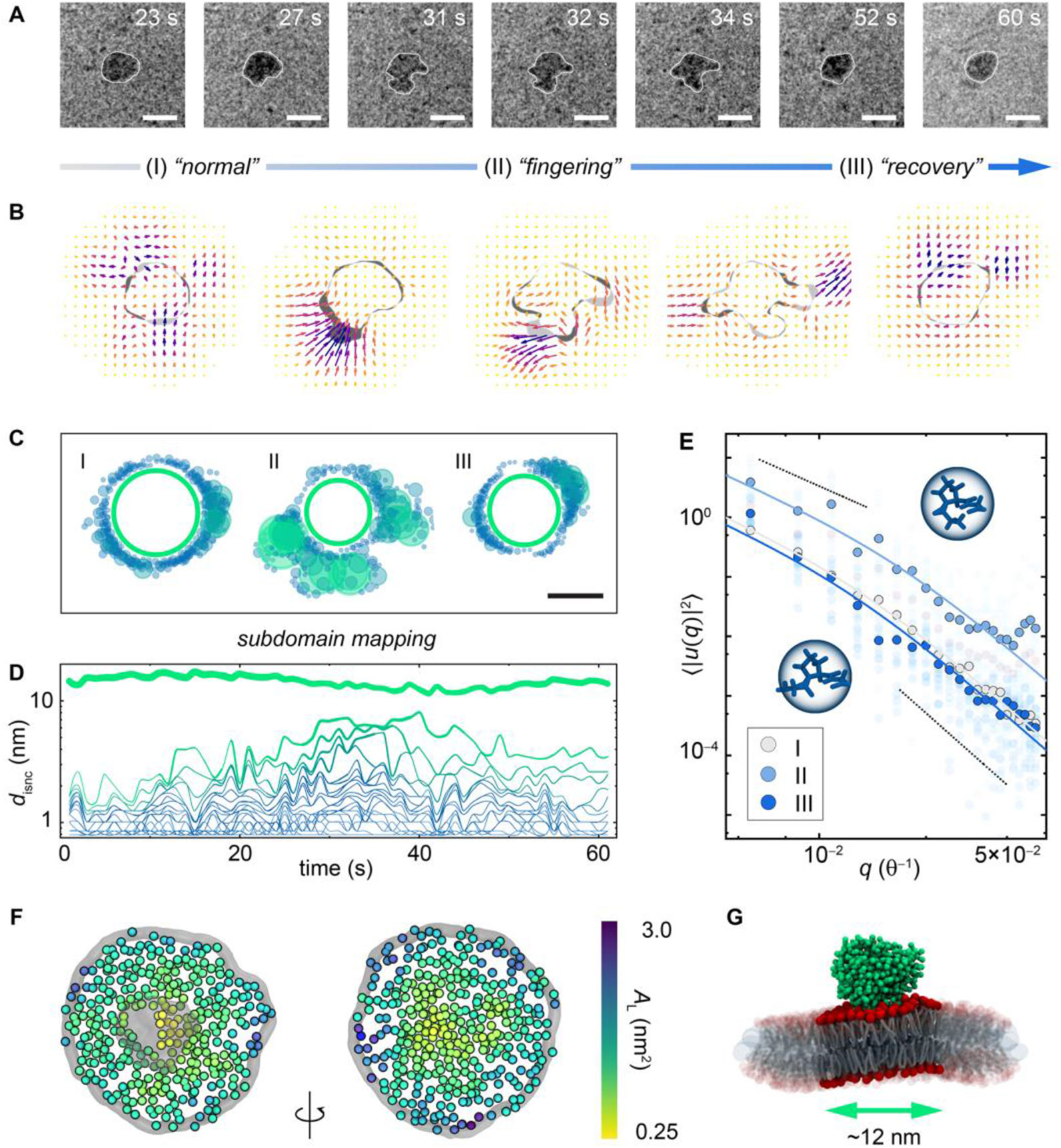
Fingering fluctuations and the role of lipid–protein interactions. (**A**) Representative snapshots of nanodiscs contorted into complex shapes, involving more drastic mass transport (**B**) and the formation of extended concave regions. The reduced visibility of CYP2J2 in these snapshots may in part be attributed to greater motion blur during vigorous fingering fluctuations. (**C, D**) Fingering converts the nanodisc from a shape that can be mostly described by a single, large domain into one that requires several zones (**Supplementary Fig. 25**; different colours denote different zones) approaching 4–5 nm in diameter. However, the largest domain consistently exceeds 12 nm in diameter. (**E**) Tension-dominated and bending-dominated fluctuation regimes are present before (I), during (II), and after (III) fingering, and the mechanics of the contour appear qualitatively recovered in (III). (**F**) Recovery processes may be facilitated by lipid–protein interactions, which lead to altered packing (area-per-lipid, *A*_L_) in the vicinity of the membrane protein, as revealed by MD simulations. (**G**) This altered packing defines a zone of the nanodisc (marked by the green arrow) that may be preserved even during extensive perturbations. Scale bars: 20 nm (**A**), 15 nm (**D**).

Recovered nanodiscs regain not only their size and shape, but also their interfacial mechanics, suggesting that fingering is related to a reversible molecular phenomenon. Specifically, while ⟨|*u*(*q*)|^2^⟩ increases by ∼20% during fingering fluctuations (**Fig. 3E**, regime II), tension- and rigidity-dominated regimes are preserved. After fingering, the mechanics of the nanodisc contour are, within experimental error, restored (**Fig. 3E**, regime I *vs*. regime III, **Supplementary Table 4**), suggesting their structural integrity. Fingering fluctuations may therefore depend on an event such as bending at the prolines that punctuate MSP helices. Note that prolines also explain why the nanodisc interface appears much more flexible than a contiguous α helix^41^. On ∼10^1^ s scales, prolines can reversibly isomerize^42^ which would change local bending angles in the MSP (**Supplementary Video 3**), and the morphology of fingering nanodiscs is consistent with the “geometry” of the proline distribution (**Supplementary Fig. 24**). Such a process is thought to occur in structural transitions of lipoprotein particles^43^, from which MSPs are derived, but has not been reported in nanodiscs, though an (unknown) extended conformational transition occurring over seconds has been suggested^44^. The large energetic barrier (∼20 kcal/mol) and time scales associated with this process may explain why fingering was not observed in MD simulations, without introducing forces to bias the system^45^. At the same time, this underscores a complementary nature of liquid-phase TEM: to detect phenomena that, to MD simulations, are rare events, but still at “single-particle” resolution.

The fact that empty nanodiscs do not recover from fingering highlights the impact of a membrane protein not only on overall nanodisc shape fluctuations (**Supplementary Figs. 10**,**15**) but also on nanodisc stability, as a consequence of lipid–protein interactions (**Supplementary Fig. 12**). Even when filled nanodiscs finger, the preserved ∼12 nm domain is large enough to harbour CYP2J2 and 2–3 nm of intervening lipids (**Fig. 3F–G**). In addition, recovery generally occurs around this region (**Fig. 3C**). MD simulations indicate that the lipids near the protein are packed more densely (**Fig. 3F, Supplementary Fig. 26**), a trait that spans to the bottom leaflet of the nanodisc that is not in contact with a membrane protein. The membrane near the protein is also compacted in the *z* direction (**Supplementary Fig. 26**). Altogether, these observations suggest that the influence of lipid–membrane protein interactions reaches not just across the few innermost annular lipid shells from where the protein is inserted, but 3–4 nm from the periphery of the protein (**Fig. 3G**). The structurally and dynamically modified zone delineated by these interactions may protect against extensive perturbations. Because empty nanodiscs lack this region, such a relatively stable domain may not exist and, given the liquid-like nature of the lipids used here, extensive fluctuations become irrevocable (**Supplementary Video 4, Supplementary Fig. 23**). While 10^1^–10^2^ nm membrane islands are believed to be present in biological membranes^27^, experimental characterization of their dynamics on such a length scale, and in particular their role in structural stability, has been challenging. We expect such insight from molecular videography can be used to understand biophysical phenomena even in macroscopic biological membrane behaviour.

## DISCUSSION

Our results show that new biophysical insight is obtained through direct, experimental observation of biomolecular dynamics in liquid at the nanometre scale. Using these fluctuations as “imaging fingerprints” of lipid–protein interactions invites future studies to understand roles of lipid composition and phase behaviour, and the effects of membrane protein structure. Of particular interest will be studies controlling the liquid chamber environment *in situ*, with temperature, electric fields, *etc*. In addition, extending our use of molecular modelling and simulation to infer submolecular-scale information from single snapshots could push the effective resolution of this molecular videography approach even further in the future. We also expect the technical and analytical approaches described here can be extended not only to a wide range of nanodiscs—with variable lipid and membrane protein composition—but also to a broad spectrum of (bio)molecular assemblies. The combination of low dose-rate imaging and graphene encapsulation appears to be critical to the success of these studies. Here, at least, we have provided evidence that biomolecules can remain sufficiently intact to present the expected shape, size, contrast, and certain fluctuations. We hope these promising, initial results encourage the substantial future work necessary to pinpoint the *specific* mechanisms whereby, for example, graphene or other “scavengers” could elicit such protective effects. Such questions present a tremendous challenge, but are rooted in important areas of basic science. Other enduring challenges are associated with our relatively low temporal resolution (ms–s) compared to many biomolecular phenomena and information lost to diffusive, rather than intramolecular, motion. Even with the advent of faster and faster electron detectors, one must begin to consider the balance between dose per frame and dose per second in terms of sample stability; though the stability limit may be defined by the dose over time, the ultimate interpretability limit may be defined by dose per image. Nevertheless, with parallel technical improvements in sample preparation and video analysis that integrate machine intelligence^46^, for example, we hope to push closer to the direct observation of proteins and other biomolecules “in action” when performing functional physical or chemical tasks, and establish a sister to cryogenic TEM in the study of molecular biophysics.

## Acknowledgments

We are grateful to Profs. Emad Tajkhorshid and Charles Schroeder of the University of Illinois for their insightful feedback on the manuscript. We also thank Drs. Giuseppe Licari and Cong Xu, formerly of the University of Illinois, for helpful discussions and guidance. This work was supported by funding from the Air Force Office of Scientific Research (grant AFOSR FA9550-20-1-0257; Q.C.), the Defense University Research Instrumentation Program (grant AFOSR FA9550-18-1-0393; Q.C.) and the National Institutes of Health (grant R01GM115584; A.D.). Experiments were carried out in the Materials Research Laboratory Central Research Facilities and Veterinary Medicine Basic Sciences Building at the University of Illinois.

## Author contributions

Conceptualization: J.W.S., A.D., Q.C.

Formal analysis: J.W.S., Q.C.

Investigation: J.W.S. (liquid-phase TEM, simulations), L.N.C. (nanodisc preparation)

Supervision: A.D., Q.C.

Visualization: J.W.S., Q.C.

Writing—original draft: J.W.S., A.D., Q.C.

Writing—review & editing: J.W.S., L.N.C., A.D., Q.C.

## References

1. Zhou, R., Huang, X., Margulis, C. J. & Berne, B. J. Hydrophobic Collapse in Multidomain Protein Folding. Science 305, 1605–1609 (2004).

2. Ball, P. Water as an Active Constituent in Cell Biology. Chem. Rev. 108, 74–108 (2008).

3. Chaplin, M. Do we underestimate the importance of water in cell biology? Nat. Rev. Mol. Cell Biol. 7, 861–866 (2006).

4. Bengtsen, T. et al. Structure and dynamics of a nanodisc by integrating NMR, SAXS and SANS experiments with molecular dynamics simulations. eLife 9, (2020).

5. Hong, M., Zhang, Y. & Hu, F. Membrane Protein Structure and Dynamics from NMR Spectroscopy. Annu. Rev. Phys. Chem. 63, 1–24 (2012).

6. Marciel, A. B., Srivastava, S., Ting, J. M. & Tirrell, M. V. Chapter Eight - SAXS methods for investigating macromolecular and self-assembled polyelectrolyte complexes. in Methods in Enzymology (ed. Keating, C. D.) vol. 646 223–259 (Academic Press, 2021).

7. Ross, F. M. Opportunities and challenges in liquid cell electron microscopy. Science 350, (2015).

8. Nakane, T. et al. Single-particle cryo-EM at atomic resolution. Nature 587, 152–156 (2020).

9. Kodera, N., Yamamoto, D., Ishikawa, R. & Ando, T. Video imaging of walking myosin V by high-speed atomic force microscopy. Nature 468, 72–76 (2010).

10. Balzarotti, F. et al. Nanometer resolution imaging and tracking of fluorescent molecules with minimal photon fluxes. Science 355, 606–612 (2017).

11. Betzig, E. et al. Imaging Intracellular Fluorescent Proteins at Nanometer Resolution. Science 313, 1642–1645 (2006).

12. Yip, K. M., Fischer, N., Paknia, E., Chari, A. & Stark, H. Atomic-resolution protein structure determination by cryo-EM. Nature 1–5 (2020) doi:10.1038/s41586-020-2833-4.

13. Shen Huaizong, Liu Dongliang, Wu Kun, Lei Jianlin, & Yan Nieng. Structures of human Nav1.7 channel in complex with auxiliary subunits and animal toxins. Science 363, 1303–1308 (2019).

14. Ye, X. et al. Single-particle mapping of nonequilibrium nanocrystal transformations. Science 354, 874–877 (2016).

15. Park, J. et al. 3D structure of individual nanocrystals in solution by electron microscopy. Science 349, 290–295 (2015).

16. Ou, Z., Wang, Z., Luo, B., Luijten, E. & Chen, Q. Kinetic pathways of crystallization at the nanoscale. Nat. Mater. 19, 450–455 (2020).

17. Ianiro, A. et al. Liquid–liquid phase separation during amphiphilic self-assembly. Nat. Chem. 11, 320–328 (2019).

18. Varano, A. C. et al. Visualizing virus particle mobility in liquid at the nanoscale. Chem. Commun. 51, 16176–16179 (2015).

19. Mirsaidov, U. M., Zheng, H., Casana, Y. & Matsudaira, P. Imaging Protein Structure in Water at 2.7 nm Resolution by Transmission Electron Microscopy. Biophys. J. 102, L15– L17 (2012).

20. Keskin, S. & de Jonge, N. Reduced Radiation Damage in Transmission Electron Microscopy of Proteins in Graphene Liquid Cells. Nano Lett. 18, 7435–7440 (2018).

21. Jonge, N. de, Peckys, D. B., Kremers, G. J. & Piston, D. W. Electron microscopy of whole cells in liquid with nanometer resolution. Proc. Natl. Acad. Sci. 106, 2159–2164 (2009).

22. Cho, H. et al. The Use of Graphene and Its Derivatives for Liquid-Phase Transmission Electron Microscopy of Radiation-Sensitive Specimens. Nano Lett. 17, 414–420 (2017).

23. Muller, M. P. et al. Characterization of Lipid–Protein Interactions and Lipid-Mediated Modulation of Membrane Protein Function through Molecular Simulation. Chem. Rev. 119, 6086–6161 (2019).

24. Brandt, E. G., Braun, A. R., Sachs, J. N., Nagle, J. F. & Edholm, O. Interpretation of fluctuation spectra in lipid bilayer simulations. Biophys. J. 100, 2104–2111 (2011).

25. Gao, Y., Cao, E., Julius, D. & Cheng, Y. TRPV1 structures in nanodiscs reveal mechanisms of ligand and lipid action. Nature 534, 347–351 (2016).

26. Denisov, I. G. & Sligar, S. G. Nanodiscs in Membrane Biochemistry and Biophysics. Chem. Rev. 117, 4669–4713 (2017).

27. Sezgin, E., Levental, I., Mayor, S. & Eggeling, C. The mystery of membrane organization: composition, regulation and roles of lipid rafts. Nat. Rev. Mol. Cell Biol. 18, 361–374 (2017).

28. Phillips, R., Ursell, T., Wiggins, P. & Sens, P. Emerging roles for lipids in shaping membrane-protein function. Nature 459, 379–385 (2009).

29. Martinez, D. et al. Lipid Internal Dynamics Probed in Nanodiscs. Chemphyschem 18, 2651–2657 (2017).

30. Li, Y., Kijac, A. Z., Sligar, S. G. & Rienstra, C. M. Structural Analysis of Nanoscale Self-Assembled Discoidal Lipid Bilayers by Solid-State NMR Spectroscopy. Biophys. J. 91, 3819–3828 (2006).

31. Sejdiu, B. I. & Tieleman, D. P. Lipid-Protein Interactions Are a Unique Property and Defining Feature of G Protein-Coupled Receptors. Biophys. J. 118, 1887–1900 (2020).

32. Shih, A. Y., Denisov, I. G., Phillips, J. C., Sligar, S. G. & Schulten, K. Molecular Dynamics Simulations of Discoidal Bilayers Assembled from Truncated Human Lipoproteins. Biophys. J. 88, 548–556 (2005).

33. Grinkova, Y. V., Denisov, I. G. & Sligar, S. G. Engineering extended membrane scaffold proteins for self-assembly of soluble nanoscale lipid bilayers. Protein Eng. Des. Sel. 23, 843–848 (2010).

34. Carnevale, L. N., Arango, A. S., Arnold, W. R., Tajkhorshid, E. & Das, A. Endocannabinoid Virodhamine Is an Endogenous Inhibitor of Human Cardiovascular CYP2J2 Epoxygenase. Biochemistry 57, 6489–6499 (2018).

35. Zhang, L., Tong, H., Garewal, M. & Ren, G. Optimized negative-staining electron microscopy for lipoprotein studies. Biochim. Biophys. Acta 1830, 2150–2159 (2013).

36. Shaw, A. W., Pureza, V. S., Sligar, S. G. & Morrissey, J. H. The Local Phospholipid Environment Modulates the Activation of Blood Clotting *. J. Biol. Chem. 282, 6556–6563 (2007).

37. Orioli, S., Henning Hansen, C. G. & Arleth, L. ıt Ab initio determination of the shape of membrane proteins in a nanodisc. Acta Crystallogr. Sect. D 77, 176–193 (2021).

38. Stone, M. B., Shelby, S. A. & Veatch, S. L. Super-Resolution Microscopy: Shedding Light on the Cellular Plasma Membrane. Chem. Rev. 117, 7457–7477 (2017).

39. Casuso, I. et al. Characterization of the motion of membrane proteins using high-speed atomic force microscopy. Nat. Nanotechnol. 7, 525–529 (2012).

40. Gan, C., Ao, M., Liu, Z. & Chen, Y. Imaging and force measurement of LDL and HDL by AFM in air and liquid. FEBS Open Bio 5, 276–282 (2015).

41. Choe, S. & Sun, S. X. The elasticity of α-helices. J. Chem. Phys. 122, 244912 (2005).

42. Osváth, S. & Gruebele, M. Proline can have opposite effects on fast and slow protein folding phases. Biophys. J. 85, 1215–1222 (2003).

43. Borhani, D. W., Rogers, D. P., Engler, J. A. & Brouillette, C. G. Crystal structure of truncated human apolipoprotein A-I suggests a lipid-bound conformation. Proc. Natl. Acad. Sci. 94, 12291 (1997).

44. Morgan, C. R. et al. Conformational transitions in the membrane scaffold protein of phospholipid bilayer nanodiscs. Mol. Cell. Proteomics MCP 10, M111.010876-M111.010876 (2011).

45. Hamelberg, D., Shen, T. & McCammon, J. A. Phosphorylation Effects on cis/trans Isomerization and the Backbone Conformation of Serine−Proline Motifs: Accelerated Molecular Dynamics Analysis. J. Am. Chem. Soc. 127, 1969–1974 (2005).

46. Yao, L., Ou, Z., Luo, B., Xu, C. & Chen, Q. Machine Learning to Reveal Nanoparticle Dynamics from Liquid-Phase TEM Videos. ACS Cent. Sci. (2020) doi:10.1021/acscentsci.0c00430.

